# Prenatal experience with language shapes the brain

**DOI:** 10.1101/2023.05.25.542259

**Authors:** Benedetta Mariani, Giorgio Nicoletti, Giacomo Barzon, María Clemencia Ortíz Barajas, Mohinish Shukla, Ramón Guevara, Samir Simon Suweis, Judit Gervain

**Affiliations:** Department of Physics and Astronomy, University of Padua, Italy; Padova Neuroscience Center, University of Padua, Italy; Department of Mathematics, University of Padua, Italy; Integrative Neuroscience and Cognition Center, CNRS and Université Paris Cité, Paris, France; Department of Developmental and Social Psychology, University of Padua, Italy

## Abstract

Human infants acquire language with striking ease compared to adults, but the neural basis of their remarkable brain plasticity for language remains little understood. Applying a scaling analysis of neural oscillations for the first time to address this question, we show that newborns’ electrophysiological activity exhibits increased long-range temporal correlations after stimulation with speech, particularly in the prenatally heard language, indicating the early emergence of brain specialization for the native language.

## Main

Human infants acquire language with amazing ease. This feat may begin early, possibly even before birth^1–5^, as hearing is operational by 24-28 weeks of gestation^6^. The intrauterine environment acts as a low-pass filter, attenuating frequencies above 600Hz^2,7^. As a result, individual speech sounds, are suppressed in the low-pass filtered prenatal speech signal, but prosody, i.e. the melody and rhythm of speech, is preserved. Fetuses already learn from this prenatal experience^5,8^: newborns prefer their mother’s voice over other female voices^1^ and show a preference for the language their mother spoke during pregnancy over other languages^3^. After birth, as infants get exposed to the full-band speech signal, they become attuned to the fine details of the sound patterns of their native language by the end of the first year of life^9–13^. What neural mechanisms allow the developing brain to learn from language experience remains, however, poorly understood. Here, we asked whether stimulation with speech may induce dynamical changes able to support learning in newborn infants’ brain activity, and whether this modulation is specific to the language heard prenatally.

We measured prenatally French-exposed newborns’ (n=49, age: 2.39 days; range 1-5 days; 19 girls) neural activity using electroencephalography (EEG) over 10 frontal, temporal and central electrode sites while infants were at rest in their hospital bassinets (Figure 1A-B). We first measured resting state activity for 3 minutes (silence 1). Then infants heard speech in three different languages, French, Spanish and English in 7-minute blocks. Finally, resting state activity was measured again for 3 minutes (silence 2; Figure 1C). The order of the languages was pseudo-randomized and counterbalanced across participants such that 17 infants heard French, 18 infants English and 14 infants Spanish as the last block prior to silence 2. Besides the prenatally heard language, French, we chose Spanish and English as unfamiliar languages to test the effects of prenatal experience. Spanish is rhythmically similar to French, while English is different^14^. Behaviorally, newborns can discriminate rhythmically different languages, even if those are unfamiliar to them, while they cannot distinguish rhythmically similar ones^15^.

**Figure 1.**
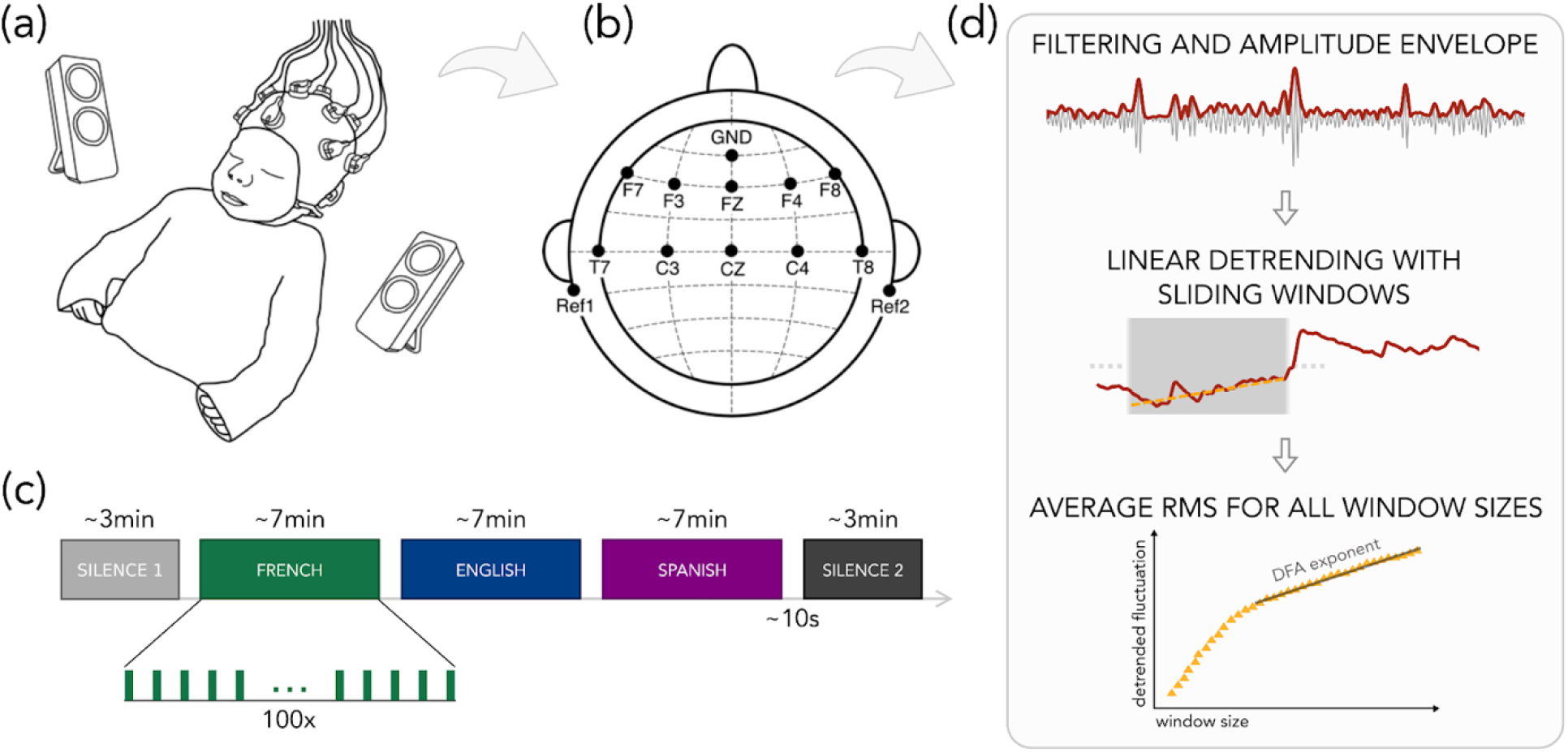
Illustration of the experimental paradigm and the analysis pipeline. (a) The experimental setup used in the study. (b) EEG channel locations. (c) The experimental design. (d) The detrended fluctuations analysis (DFA) analysis.

By comparing resting states before and after linguistic stimulation, this paradigm allowed us to address two questions. First, whether language exposure impacts neural dynamics in the infant brain: plastic changes *immediately* following exposure to speech may underlie infants’ ability to learn about the sound patterns they hear^16,17^. We were not interested in the newborn brain’s responses to different languages—a very interesting question, which has received ample attention in the literature^18–22^, including using data from this study^23^. Rather, we asked whether exposure to speech makes lasting changes in neural dynamics, supporting learning and memory.

Second, by testing whether these plastic changes occur after exposure to all languages or only after the language heard prenatally, we asked whether prenatal experience already shapes neural circuitry. If prenatal experience already plays a role, then newborns may show greater plastic changes after exposure to the language heard prenatally than after unfamiliar languages. As an especially stringent test, we compared the native language not to one, but to two unfamiliar languages, including a rhythmically similar language, which newborns *cannot* discriminate from the native language behaviorally^15^.

We expected changes in neural dynamics specifically at low frequencies, as the prenatal speech signal is low-pass filtered and mainly consists of low-frequency information, i.e. prosody^2,7,24^. According to the embedded neural oscillations model^25,26^, oscillations between 1-3 Hz, i.e. the delta band, underlie the processing of large prosodic units, such as utterances and phrases, oscillations between 4-8 Hz, i.e. the theta band, underlie the processing of syllables, while oscillations above 35 Hz, i.e. the gamma band, are related to the processing of phonemes. We thus predicted that prenatal effects will specifically target the delta and theta bands, as linguistic units corresponding to those frequency bands are the ones present in the prenatal signal.

To assess changes in neural dynamics induced by speech, we conducted two analyses: (i) detrended fluctuation analysis (DFA) and (ii) the autocorrelation between the fluctuations of electrophysiological activity before and after stimulation.

Detrended fluctuation analysis^27^ (DFA) is a robust method for measuring the statistical self-similarity and temporal correlation of time series data. Specifically, DFA measures the scaling properties of signals, quantified by the strength of long-range temporal correlations (LRTC). The strength of LRTC is given by the scaling exponent α, i.e. the exponent of the power law relation between the average fluctuations of the signal and different time scales (widow sizes). DFA has been successfully applied to distinguish between healthy and pathological neural activity^28,29^ and to identify the onset of sleep^30^.

The scaling exponent, α, is an estimate of the Hurst parameter, which indicates whether a process generating a given time-series has ‘memory’ in the sense that the state at a given moment does or does not relate to prior states. A value of α=0.5 indicates an uncorrelated process, with the current state having no dependence on prior states, while α<0.5 indicates an anti-correlated process, with previous states being avoided, leading to smaller fluctuations at larger timescales than would be expected by chance. Scaling exponents 0.5>α>1 indicate self-affine stationary processes with positively correlated memory, i.e. whereby the system is more likely to enter states that it had visited before. Human adult EEG typically exhibits Hurst exponents of ∼0.70 ^31^, i.e. is a self-affine, positively correlated process.

Here, we hypothesize that newborn brain processes, as revealed by EEG, show evidence of language learning, i.e. lasting changes in brain dynamics after exposure to language. Specifically, upon encountering a learned, i.e. previously already activated brain state triggered by previous language experience, newborns’ brain processes maintain these triggered states, resulting in an increase in the value of α. Further, such an increase would be expected specifically for the prenatally heard, i.e. previously experienced language, and not for unfamiliar languages, since the brain states they trigger would not be similarly privileged.

To calculate DFA, EEG recordings from the resting state periods were pre-processed using standard pipelines for infant auditory EEG data^32–34^. Subsequently, signals were band-passed between 4-8 Hz to obtain theta oscillations and between 30-60 Hz to obtain gamma oscillations. (The delta band could not be included in the analysis, as the 3-minute duration of the resting state periods did not provide sufficient data for the calculations.) We then extracted the amplitude envelope of the oscillations and performed DFA^27,35^. For each channel of each participant, we selected window sizes equally spaced on a logarithmic scale. For each size, we split the signals into windows with 50% overlap, detrended each window through a least-squares fit, and calculated the standard deviation of the detrended signal. We then obtained the fluctuation function as the average standard deviation of the detrended signal computed over the windows, as a function of window size. By plotting this on a log-log scale, the scaling exponent was obtained using a linear fit (see Methods).

As the scaling distributions show (Figure 2B-F), oscillations exhibited power law scaling on time scales between 0.86-20sec in theta, and between 0.43-20sec in gamma. To test whether LRTCs get stronger after stimulation with language, we applied a linear mixed effects model to the DFA exponents with Resting State Period (silence 1 / silence 2) as a fixed factor and Participants as a random factor (see Methods for model selection). In theta, scaling exponents showed a statistically significant increase from silence 1 (α=0.76±0.06) to silence 2 (α=0.81±0.08; slope=0.05, p=0.0005). By contrast, in gamma, a significant decrease was observed (silence 1: α=0.95±0.1269; silence 2: α=0.88±0.16; slope= -0.07, p=0.01).

**Figure 2.**
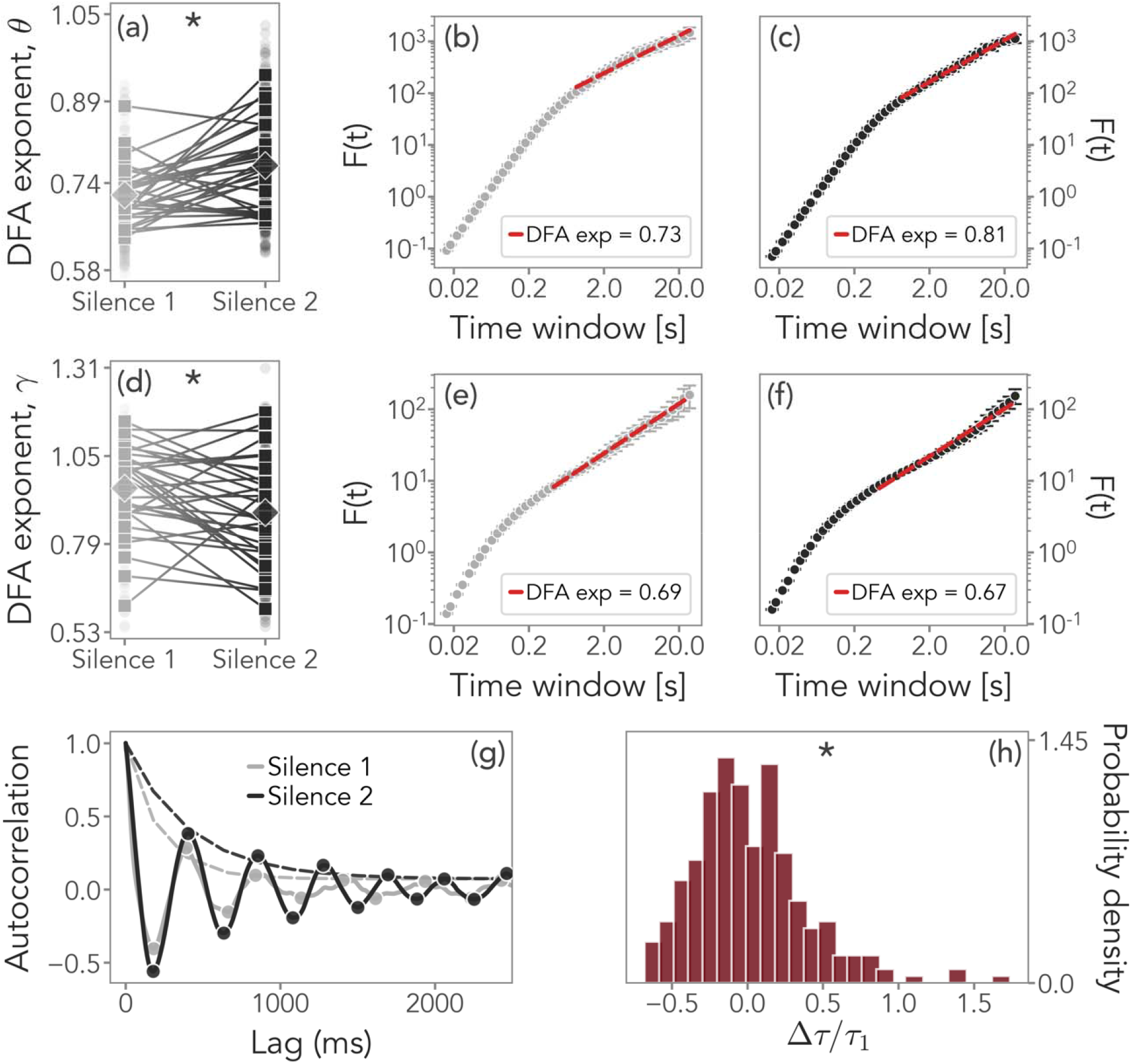
DFA exponents and temporal correlations. (a-c) Comparison of the average DFA exponents in the theta band during silence 1 (gray) and silence 2 (black) in channel F8 as an example. Squares indicate individual DFA exponents, diamonds the average. Statistically significant results are indicated by an asterisk. (d-f) DFA exponents in the gamma band. (g) Comparison of the autocorrelation function in channel F8 during silence 1 (gray) and silence 2 (black) and the fits of the corresponding exponential envelopes (dashed lines) whose decay defines the autocorrelation time. (h) Distribution of the relative change in the autocorrelation times, obtained for all channels and all subjects. Its significantly positive skewness suggests that stronger temporal correlations are present after language stimulation.

These results confirm our hypothesis and have two implications. First, they reveal for the first time that exposure to speech rapidly increases long-range correlations in neural activity, thereby highlighting how language experience may shape the brain and contribute to learning. As in adults^31^, newborns’ electrophysiological activity showed the properties of a self-affine stationary process with positively correlated memory, with exponents ∼0.7-0.8 before stimulation. Importantly, long-term correlations got strengthened after stimulation for several minutes, providing evidence for learning. Second, LRTCs were enhanced specifically in the theta band, associated with the syllabic rate, i.e. speech units experienced in utero, as predicted, but not in gamma. The gamma band actually showed a slight decrease in long-range correlations. This increase in neural resources in theta as compared to gamma oscillations may be related to the greater importance of prosodic units such as syllables in infants’ representation of speech at birth due to their prevalence in the prenatal signal, as suggested by behavioral results^36–38^ and theoretical models^39^.

To test the role of prenatal experience directly, we compared changes in LRTCs as a function of the last language heard before silence 2 (Figure 3). A linear mixed effects model with Resting State Period (silence 1 / silence 2) as a fixed factor and Participants as a random factor showed that in the theta band, only infants who listened to French last showed a significant increase in scaling exponents from silence 1 (α=0.76±0.05) to silence 2 (α=0.82±0.07; slope=0.065, p=0.0003). Those exposed to Spanish (slope=0.05, n.s.) or English (slope=0.04, n.s.) last did not. In the gamma band, no change was observed for newborns exposed last to French (slope= -0.04, n.s.) or Spanish (slope= -0.01, n.s.), while a significant decrease from silence 1 (α=0.99±0.08) to silence 2 (α=0.86±0.14) was found for those exposed last to English (slope= -0.14, p=0.003).

**Figure 3.**
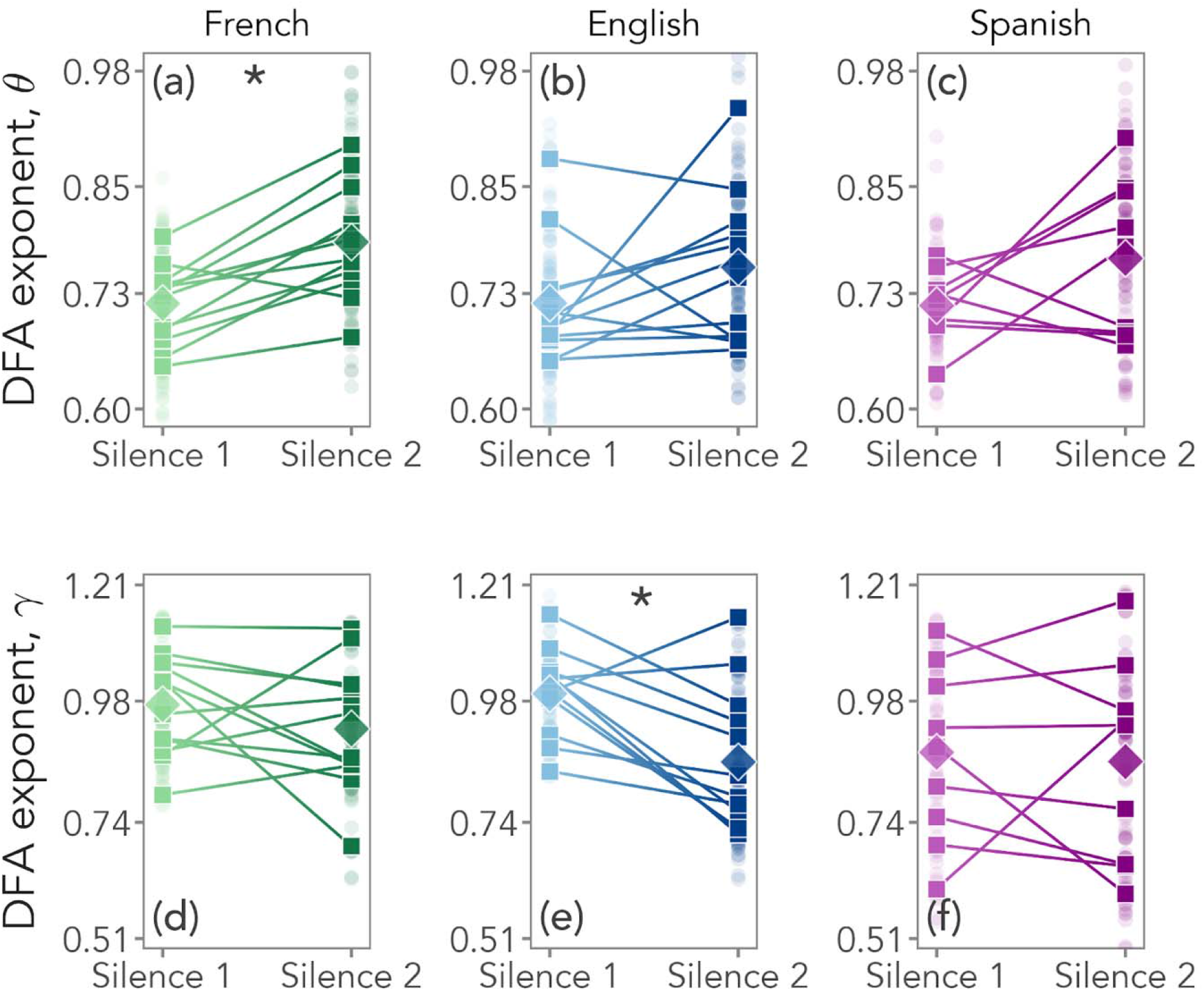
DFA exponents as a function of the language heard last before silence 2. (a-c) DFA exponents in the theta band for participants who heard French (green), English (blue), and Spanish (violet) as the last language before silence 2. (d-f) DFA exponents in the gamma band.

As a second approach, we also estimated the autocorrelation function of the fluctuations of the signal’s temporal correlations. We found that autocorrelations exhibited an oscillatory decay (Figure 2G), and we computed the corresponding autocorrelation time by fitting their exponential envelope (see Methods). We thus compared temporal correlations before and after language stimulation. The positive skewness (1.05, p<0.00001) and kurtosis (2.19, p=0.00002) of the distribution of the relative change in autocorrelation times (Figure 2H) suggest that language stimulation indeed leads to increased temporal correlations and therefore to sustained activity.

Taken together, these results provide the most compelling evidence to date that language experience already shapes the functional organization of the infant brain, even before birth. Exposure to speech leads to rapid, but lasting changes in neural dynamics, enhancing long-range temporal correlations and thereby increasing infants’ sensitivity to previously heard stimuli. This facilitatory effect is specifically present for the language and the frequency band experienced prenatally.

From a broader perspective, our findings document, for the first time, power law scaling of neural activity during language processing in the newborn brain. This statistical property is a hallmark of critical phenomena, and it has been suggested that criticality in the brain is linked to states of optimal information transmission and storage^40–42^. The newborn brain may thus already be in an optimal state for the efficient processing of speech and language, underpinning human infants’ surprising language learning abilities.

## Methods

The EEG data from this study was acquired as part of a larger project that aimed to investigate speech perception and processing during the first two years of life^23^.

### Participants

The protocol for this study was approved by the CER Paris Descartes ethics committee, Université Paris Descartes, Paris, France (currently, Université Paris Cité). All parents gave written informed consent prior to participation, and were present during the testing session. We recruited participants at the maternity ward of the Robert Debré Hospital in Paris, and we tested them during their hospital stay. The inclusion criteria were: i) being born full-term and healthy, ii) having a birth weight > 2800 g, iii) having an Apgar score > 8 at 5 and 10 minutes after birth, iv) being maximum 5 days old, and v) being born to French native speaker mothers who spoke French at least 80% of the time during the last trimester of the pregnancy according to self-report. We recruited a total of 54 newborns, and excluded 5 participants due to fussiness and crying (n = 4) or technical problems (n = 1), resulting in 49 newborns who completed the study. Of these, we excluded 16 participants due to bad data quality in the silence 1 or silence 2 periods (see Data Analysis for data quality criteria). Thus, electrophysiological data from 33 newborns (age 2.55 ± 1.33 d; range 1-5 d; 16 girls, 17 boys) were included in the final analysis. Among these participants, 12 infants heard French as the last stimulation block, while 12 heard English, and 9 heard Spanish.

### Stimuli

We tested infants in five blocks: at rest (silence 1), during three blocks of speech stimulation with three different languages and again at rest (silence 2; Figure 1C). While at rest, no stimulus was presented. During speech stimulation, the following three languages were presented in separate blocks: the infants’ native language (French), a rhythmically similar unfamiliar language (Spanish), and a rhythmically different unfamiliar language (English). Stimuli consisted of sentences taken from the story “Goldilocks and the Three Bears” and its translation equivalents in the other two languages. Three sets of sentences were used, where each set comprised the translation of a single utterance into the 3 languages (English, French and Spanish). The translations were created so as to match sentence duration across languages within the same set both in number of syllables and in absolute duration. All sentences were recorded in mild infant-directed speech by a female native speaker of each language (a different speaker for each language), at a sampling rate of 44.1 kHz. There were no significant differences between the sentences in the three languages in terms of minimum and maximum pitch, pitch range and average pitch. The intensity of all recordings was adjusted to 77 dB. Further details of the stimuli are available in^23^.

### Procedure

Newborns were tested in a dimmed, quiet room at the Robert Debré Hospital in Paris. During the recording session, newborns were comfortably at rest in their hospital bassinets (Figure 1AB). The speech stimuli were delivered bilaterally through two loudspeakers positioned on each side of the bassinet using the experimental software E-Prime. The sound volume was set to a comfortable conversational level (∼65-70 dB). We presented participants with one sentence per language, repeated 100 times. The experiment consisted of 5 blocks: one initial resting state block (silence 1), three language blocks, and one final resting state block (silence 2; Figure 1C). Each resting state block lasted 3 minutes, while each language block lasted around 7 minutes. An interstimulus interval of random duration (between 1 – 1.5 s) was introduced between sentence repetitions, and an interblock interval of 10 s was introduced between language blocks (Figure 1C). The order of the languages was pseudo-randomized and counterbalanced across participants. The entire recording session lasted about 27 minutes.

### Data acquisition

We recorded EEG data with active electrodes, using a Brain Products actiCAP & actiCHamp acquisition system (Brain Products GmbH, Gilching, Germany). We used a 10-channel layout to acquire cortical responses from the following scalp positions: F7, F3, FZ, F4, F8, T7, C3, CZ, C4, T8 (Figure 1B). We chose these recording locations, because this is where auditory and speech perception related neural responses are typically observed in newborns and young infants^33,34^ (channels T7 and T8 were previously labelled T3 and T4, respectively). We used two electrodes placed on each mastoid for online reference, and a ground electrode placed on the forehead. Data were referenced online to the average of the two mastoid channels. Data were recorded at a sampling rate of 500 Hz, and filtered online with a high cutoff filter at 100Hz, a low cutoff filter at 0.1Hz and an 8 kHz (−3 dB) anti-aliasing filter. The electrode impedances were kept below 140 kΩ.

### Data Analysis

We pre-processed and analyzed the data using custom Python and R scripts.

#### Pre-processing

First, we filtered the raw EEG signals with a 50Hz notch filter to eliminate the power line noise. Then, we band-pass filtered the denoised EEG signals between 1 and 100 Hz using a zero phase-shift Chebyshev filter. Then, we submitted the recordings to a rejection process to exclude noisy and artifact-contaminated data. First, we identified channels with amplitudes exceeding ± 200μV. If that happened in the first or last 30 seconds of the 3min resting periods, we rejected only the compromised data segment. Otherwise, we rejected the whole channel. This way, we ensured that we had at least 150 seconds of continuous artifact-free recordings, as needed for the DFA analysis. We then rejected channels whose power spectrum (in dB) was above or below the participant’s average power spectrum by more than 4,5*10^−5^. Finally, the procedure was validated by visual inspection to remove any residual artifacts. Participants who had fewer than 5 valid channels after channel rejection were not included in the data analysis (n=15). On average, participants contributed 9 clean channels out of the 10 total channels (SD 1.5 for silence 1,and 0.9 for silence 2). All data pre-processing and rejection was conducted in batch using the above defined criteria, prior to statistical analysis.

#### DFA analysis

Detrended fluctuation analysis (DFA) was performed on the silence 1 and silence 2 blocks.

The signal was filtered in the theta (4-8 Hz) and low gamma (30-60 Hz) bands. The delta band (1-3 Hz), although theoretically relevant, could not be included in the analysis: the 3-minute duration of the resting state periods did not provide sufficient data for fitting the estimate of the DFA exponent in this slow band.

The signals were filtered with a FIR (Finite Impulse Response) filter, following standard practice^27^, since it avoids introducing artifactual long range correlations, unlike other filtering procedures. The order of the filter was varied across bands so as to cover at least two cycles of the lowest frequency of the band, and was thus set to 2*(sampling frequency/lowest frequency).

DFA analysis was then performed on the amplitude envelope of the filtered signals, i. e. on the amplitude extracted from the analytic representation of the signal, with the following steps^27,35^. The cumulative sum of the time series was computed to create the signal profile. A set of window sizes, T, which were equally spaced on a base 2-logarithmic scale between the lower bound of eight samples and the length of the signal, was selected. For each window length t ∈ T, the signal profile was split into a set W of separate time series of length t, which have 50% overlap. For each window w ∈ W the linear trend was removed (using a least-squares fit) from the time series to create the signal w_detrend. The standard deviation of the detrended signal σ(w_detrend) was calculated in each window. Then, the fluctuation function F was calculated as the mean standard deviation of the detrended signals over all identically sized windows, as a function of the window size: F(t) = mean(σ(w_detrend(t))). The fluctuation function was plotted for all window sizes, T, on logarithmic scales. The DFA exponent, α, is the slope of the trend line in the range of time-scales of interest and can be estimated using linear regression. The lower bound for the linear regression fit was chosen by comparing the fluctuation function of the signal with a white noise surrogate, filtered using the same filter as the data. The lower bound was placed where the expected white noise scaling (α = 0.5) started to appear^27^. This lower bound was a function of the band chosen, i.e. the lower the frequency band, the higher the bound, as expected. The higher bound was set at 20 seconds, i.e. approximately 15% of the minimum signal length, i.e. 150 seconds.

The DFA exponent can be interpreted as an estimate of the Hurst parameter^27^, i.e. the time series is uncorrelated if α = 0.5. If 0.5 < α < 1, then there are positive correlations present in the time series, i.e. the series shows larger fluctuations on longer time-scales than expected by chance. If 0 < α < 0.5, then the time series is anti-correlated, i.e. fluctuations are smaller in larger time windows than expected by chance.

#### Statistical analysis

Two sets of statistical analyses were conducted: one to assess the overall impact of speech stimulation on neural dynamics, and one to assess whether speech in different languages had different impact. For the first analysis, the DFA exponents were entered into a linear mixed effects models to test whether there is a significant difference in the DFA exponents of the signals before and after speech stimulation. We built and compared all the possible models. Model selection was based on the Akaike Information Criterion (AIC)^43^. The models were implemented using the *lme4* and *lmertest* packages in R^44^, and the parameters were estimated optimizing the log-likelihood criterion. In both the theta and the gamma bands, the best fitting model included Resting State Period (silence 1 / silence 2) as a fixed factor and Participant as random factor.

For the second analysis, linear mixed effects models were run, dividing the subjects in three groups on the basis of the last language heard prior to silence 2. For participants who heard French last, the best fitting model in the theta band included Resting State Period (silence 1 / silence 2) as a fixed factor and Participant as a random factor. The same model was also the best in the gamma band for participants who heard English last. For all other comparisons, the best fitting model only included Participant as a random factor.

#### Autocorrelation times

The connected autocorrelation function *c*_*i*_(*t*) of the i-th channel of the EEG signal is defined as *c*_*i*_(*T*) = ⟨*x*_*i*_(*t*_0_) *x*_*i*_(*t*_0_ + *t*)⟩ − ⟨*x*_*i*_(*t*_0_)⟩ ⟨*x*_*i*_(*t*_0_ + *t*)⟨, where ⟨ ⟩ denotes an average over all times t0 and xi(t) is the signal of the i-th channel. *c*_*i*_(*t*) describes the temporal correlations of the fluctuations of *x*_*i*_(*t*). We found that the autocorrelation function *c*_*i*_(*t*) typically displays damped oscillations, decaying with an exponential envelope *c*_*i*_(*t*) ∝ exp(−*t/τ*_*i*_), with *τ*_*i*_ being the autocorrelation time.

We estimated, for the i-th channel of the n-th subject, the autocorrelation time in silence 1, 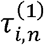, and silence 2, 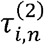, by fitting the exponential envelope of *c*_*i,n*_(*t*) with standard maximum likelihood methods. We then computed the relative change in autocorrelation time 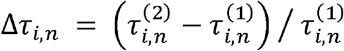, the corresponding EEG activity displays enhanced temporal correlations. To assess whether autocorrelation times increase on average across channels and subjects, we estimated the empirical probability distribution of the relative change *p*(*s*) = ∑_*i,n*_ *δ*[*s* − Δ *τ*_*i,n*_], where δ is the Dirac delta function. Positive skewness suggests that activity in silence 2 has larger correlation times than silence 1.

## Data Availability

The data that support the findings of this study are being made available at https://osf.io/g2rvc/

## Code Availability

The code to analyze the data and generate all figures of this manuscript is available at https://osf.io/g2rvc/

## Acknowledgements

The study was funded by an ERC Consolidator Grant “BabyRhythm” nr. 773202 to JG and a FARE grant nr. R204MPRHKE from the Italian Ministry for Universities and Research to JG and SSS.

## Author Information

### Contributions

JG and SSS conceptualized the study, MCOB performed the experiment, BM, GN, GB, RG performed the analyses, JG and SSS supervised the study and the analyses, JG and SSS secured funding, all authors contributed to the first draft, all authors edited and reviewed the manuscript.

## Supplementary material

**<u>Supplementary materials.docx</u>**

